# Pollen DNA reveals urban cavity-nesting bees feed primarily on weeds and frequently nest in urban farms

**DOI:** 10.1101/2025.03.09.642154

**Authors:** Katherine J. Turo, Rodney T. Richardson, Molly Frabotta, Reed M. Johnson, Mary M. Gardiner

**Affiliations:** Department of Entomology, The Ohio State University, Columbus, OH, USA; Department of Ecology, Evolution, and Natural Resources, Rutgers University, New Brunswick, NJ, USA; Department of Entomology, The Ohio State University, Wooster, OH, USA

**Keywords:** pollen metabarcoding, urban greenspaces, landscape connectivity, native wildflower, pollinator foraging

## Abstract

Cities have been acclaimed as hotspots for bee biodiversity and potential conservation targets, leading to continued investment in urban pollinator plantings. However, newly created habitats are rarely assessed for their efficacy in supporting bee fitness or the extent to which bees use seeded wildflowers. We compared urban bee nesting in targeted “pocket prairie” pollinator plantings versus urban farms that were intended to support multiple ecosystem services in Cleveland, Ohio, USA. We used trap nests to evaluate nesting success of cavity nesting bees and pollen metabarcoding to determine whether bees collected pollen from seeded plantings during nest provisioning. Pollen DNA revealed most bee-collected pollen was from urban spontaneous vegetation (or “weeds”) in Fabaceae, especially *Trifolium* spp. We also found that urban farms harbored more native bee larvae than targeted pollinator plantings. Finally, when bee nests were situated in a landscape with greater greenspace connectivity, we observed more native bee larvae and greater plant diversity in bees’ nesting provisions. Collectively, these findings suggest that multi-service greenspaces like urban farms can provide important urban pollinator habitat, and greenspace value for bees is driven by resident weeds and greenspace configuration.

**Open Research Statement:** Data are not yet provided. Upon acceptance data will be archived in Dryad Digital Repository. Each applicable R package is cited in the text; no novel code is presented. Larval DNA sequences will be available through BOLD, Barcode of Life Data Systems; Pollen DNA reads will be available through Dryad; Raw ecological data (larvae abundance, bloom characteristics, landscape structure) will be available through Dryad.

**Highlights:**

- Exotic weeds are cavity nesting bees’ dominant pollen source in cities
- Pollen DNA improves foraging observations across a diverse urban matrix
- Native bee reproduction was greater in urban farms than pollinator plantings
- Multi-service urban farms can promote win-win conservation scenarios
- Greenspace connectivity modulates bee foraging and nesting success

## Introduction

Scientists, policymakers, and lay people are commonly invested in supporting urban pollinators (Dicks et al., 2016; Hall et al., 2017), but their motivations can be distinct and apply to different subsets of the urban bee community. First, cities often contain diverse bee communities and even rare bee species (Baldock et al., 2015; Senapathi et al., 2017; Sirohi et al., 2015), leading to increased conservation interest in maintaining native bee biodiversity in cities or facilitating new species filtering into the urban environment from regional species pools (Aronson et al., 2016, Pham et al. *in revision)*. A separate, but related, motivation for supporting urban bee populations is the provision of pollination services to urban farms and community gardens. There are roughly 800 million urban farmers worldwide (FAO) who stand to benefit from enhanced pollination, and wild bees are documented as enhancing crop quality and quantity in cities (Lowenstein et al., 2015; A. Potter & LeBuhn, 2015). A key difference between these motivations is that conservation projects focus on native biodiversity while delivery of pollination services can be achieved by supporting dominant bees (Kleijn et al., 2015), including exotic species, which are highly abundant in urban environments (Wenzel et al. 2020).

For both conservation and pollination service goals, the main method typically used to support wild bee populations in cities is the creation of bee habitat in the form of “pollinator plantings”, i.e., urban greenspaces seeded or planted with flowering forbs (Turo & Gardiner, 2019). Additional forage is thought to support bees as floral abundance strongly predicts bee abundance and diversity, even across anthropogenically disturbed landscapes (Hyjazie & Sargent, 2022; Winfree et al., 2011). Likewise, increased floral richness is thought to improve bee nutrition as richer floral resources allow bees collect pollen from different plant species, balancing their nutritional requirements (Vaudo et al., 2020) and leading to higher nutritional quality (Trinkl et al., 2020). Often, pollinator plantings focus on seeding native plants, as native species sometimes attract richer bee communities (Matteson & Langellotto, 2011; Pardee & Philpott, 2014, Pham et al. *in revision*) and can facilitate specialist foragers persisting in cities (Casanelles-Abella et al., 2022; Hanley et al., 2014). In addition, pollinator plantings can provide semi-natural habitat for nesting within the urban matrix and have been associated with greater nesting productivity for cavity nesting bees (Turo & Gardiner, 2021).

Despite widespread recommendations of pollinator plantings, there are few guidelines for how to design urban bee habitat and assessments of plantings’ efficacy are rare (Hicks et al. 2016; Pham et al. *in revision*). When plantings have been assessed, there is mixed evidence of plantings benefit to bees, and the size of a pollinator planting affects its use by bees (Blackmore & Goulson, 2014; Matteson & Langellotto, 2011; Simao et al., 2018). Evidence of plantings’ positive impact on bee populations comes from observations of bee foraging, where both native and exotic bees are documented visiting seeded plants (Blackmore & Goulson, 2014; Turo et al., 2021). But, cities are already intensively cultivated (Grimm et al., 2008), and contain high abundances of naturally established weeds (“urban spontaneous vegetation”, Riley et al. 2018) and ornamental flowers (Goddard et al., 2010), that bees are known to feed on (Lowenstein et al., 2019; Potter et al., 2019; Turo et al., 2021). To date, it is still unclear to what extent bees forage on flowers seeded in pollinator plantings versus adjacent residential yards and public greenspaces (e.g., parks, community gardens) (but see—Potter et al., 2019). Likewise, there is limited evidence whether use of an urban pollinator planting translates to improved bee fitness as most studies focus on community metrics like abundance and diversity (Brant et al., 2022), which may not reflect bee health or population growth. In addition, it is also possible that multi-service greenspaces that are not targeted towards bee conservation, like urban farms or gardens, perform equally well or better as habitat for urban bees (Felderhoff et al., 2023; Sivakoff et al., 2018). However, a direct comparison of large-scale pollinator plantings to other multi-service greenspaces in an urban environment is lacking.

Finally, bee behavior is shaped by urban landscapes, so it is critical to consider how spatial context can modulate different urban habitat’s value to bees. For example, as central place foragers, bees are predicted to optimize their foraging behavior by avoiding smaller, more isolated patches (Harrison & Winfree, 2015), and thus may not visit a high-quality urban greenspace if its’ location is suboptimal. Certainly, studies have confirmed that urban fragmentation can impede bee movement and limit foraging distance, even for highly mobile bumble bees (Bhattacharya et al., 2003; Van Rossum & Triest, 2012). Similarly, increasing proportions of impervious cover are also associated with declines in urban pollination services (Bennett & Lovell, 2019). And, greenspace configuration across an urban landscape can affect bee nesting patterns (Casanelles-Abella et al., 2022; Turo & Gardiner, 2021) and dictate which bees are able to assemble in a habitat, i.e., bee abundance, species richness, nesting guild, and body size (Marini et al., 2014; Sivakoff et al., 2018; Wenzel et al., 2020). Thus, although placement of new pollinator plantings is often dictated by sociological factors (Turo & Gardiner, 2019), landscape context is a critical ecological concern that needs to be considered when assessing the value of different urban greenspaces to bees and guiding future efforts.

As custom pollinator plantings continue to be installed in cities, it is important to clarify what benefits we can expect from seeded native plantings, and how these benefits compare to multi-service greenspaces that are not specifically targeted to bee conservation but may nonetheless provide foraging and nesting habitat. In this study, we evaluated urban bee nesting and pollen collection across a large network of native plantings and multi-service urban farms in Cleveland, Ohio, USA. We asked:

(1) Do urban bees use seeded native plants for nest provisioning?
(2) Does urban bee pollen collection vary with species identity and nest location?
(3) Is native bee fitness enhanced by targeted plantings or their configuration within the landscape?

Here, we focus on cavity nesting bees, which are solitary pollinators that nest in bored holes, reeds, or mud tubes. Cavity nesters are the predominant nesting niche that comprises many urban bee communities (Wenzel et al., 2020). Because of their unique nesting ecology, we were able to recover whole pollen provisions that maternal cavity nesting bees collected for their young across multiple foraging trips, and subsequently use DNA barcoding to identify larval species and their pollen provisions. Pollen metabarcoding is a recent, innovative technique that enables quantification and identification of plant DNA from an aggregate sample of pollen (Bell et al., 2023). A benefit of this technique is that we can recover foraging records that would be difficult to observe in a city because of the high diversity and abundance of flowering plants as well as private and public regulations that often restrict access to land (Sponsler et al., 2020). Recent studies have employed pollen metabarcoding to characterize honey bee foraging in urban environments (Richardson et al., 2021; Sponsler et al., 2020; Tanaka et al., 2020). But, to date, only a few studies have investigated pollen collection by wild urban bees (Casanelles-Abella et al., 2022; Dürrbaum et al., 2023; Potter et al., 2019), and to the best of our knowledge, none use pollen metabarcoding to understand bee foraging in North American cities.

## Methods

### Field data collection

Our study took place in the post-industrial, legacy city of Cleveland, Ohio, USA. Cleveland is one of 350 legacy cities worldwide (Rieniets, 2009), which are characterized by population decline and abundant vacant land following a sustained economic downturn and the removal of urban infrastructure (Haase et al., 2014). One result of this is that Cleveland hosts around 1,600+ hectares of vacant land. As part of a larger Cleveland Pocket Prairie Project (Turo et al., 2021), we transformed sixty-four vacant lots into native wildflower habitats, or urban “pocket prairies”, across Cleveland in a large-scale, manipulated field experiment. For this study, we collected nesting bees and their pollen provisions from a subset of ten urban prairies as well as nine pre-existing urban farms (Figure 1).

**Figure 1.**
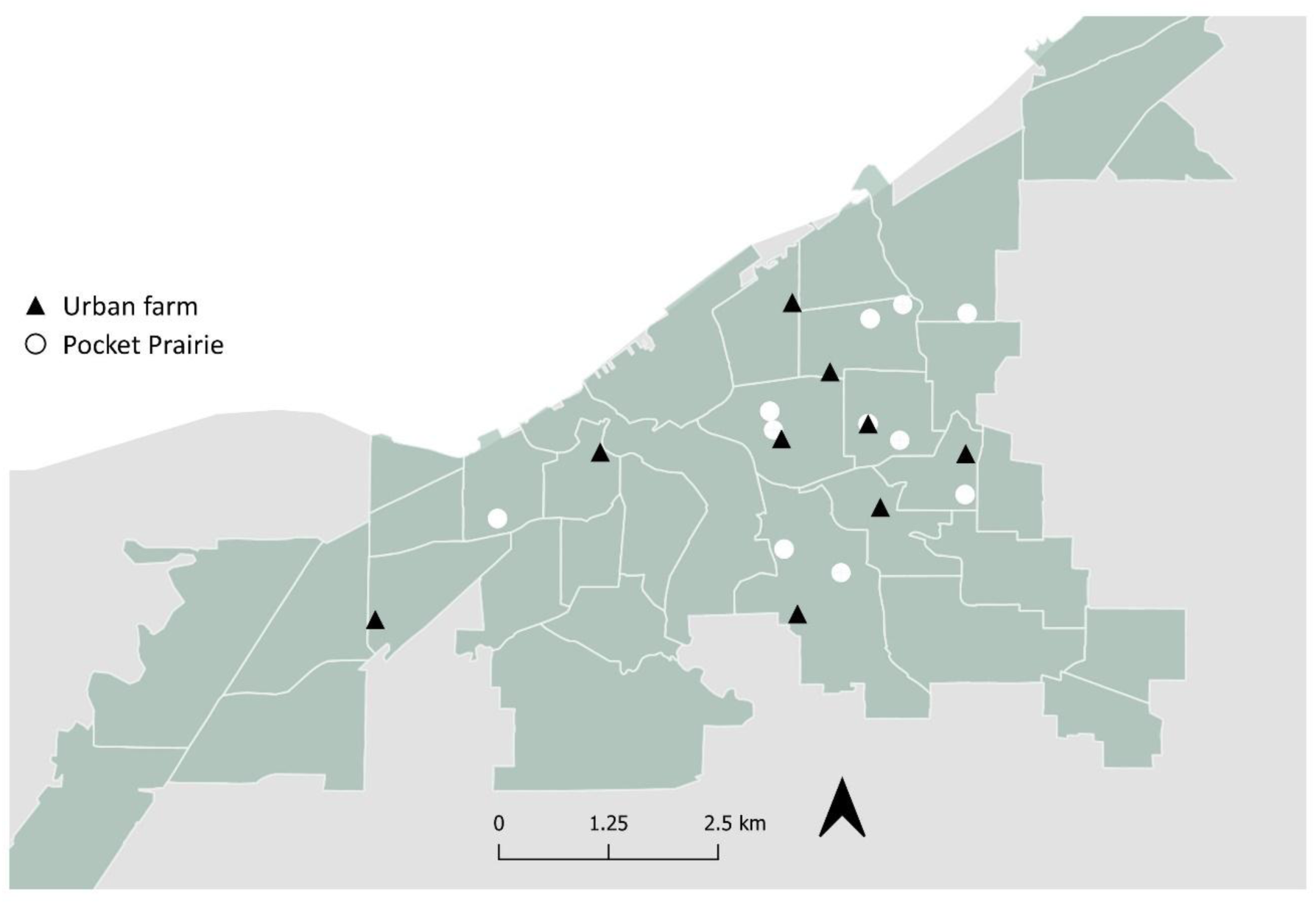
Urban greenspace study system in Cleveland, Ohio. We sampled nine urban farms and ten “pocket prairie” conservation greenspaces spread across six neighborhoods in Cleveland, Ohio, USA. Prairies were 30 x 12 m vacant lots that were seeded with native forbs three years prior to this study (Appendix 1). Urban farms focused on vegetable production and varied in size from 2.43 ha to 0.10 ha. Farms were also established on previously vacant land and were managed privately by several organizations: Bay Branch Farm, Cleveland Botanical Gardens, Cuyahoga County Extension, Kinsman Farm, and Ohio City Farm.

Urban prairies were 30 x 12 m (0.03 ha) plots of lands that were established in 2014 as insect conservation habitat. Prior to urban prairie establishment, the vacant lots were seeded with turf grass and occupied by urban spontaneous vegetation (Riley et al., 2018). During establishment, urban prairies were treated twice with herbicide and then seeded with a broadcast mixture of 3 prairie grasses and either 8 (n=4) or 16 (n=6) native forbs (Appendix 1). Overseed mixtures of an additional 4 or 6 native forbs species were added in 2016 to improve site appearance during plant establishment. Similar to the urban prairies, urban farms were developed on formerly vacant land and varied in size from 0.10 ha to 2.43 ha. All farms focused on vegetable production but were also frequently seeded with ornamental flowers in field margins or in raised beds.

At each greenspace, we collected cavity nesting bees with trap nests consisting of 30 cardboard nesting straws of 4 mm, 6 mm, and 8 mm diameter (Crown Bees, Woodinville, WA). Trap nests were deployed for two years (2017, 2018) from the early spring to summer (April 1^st^ to July 31^st^). Importantly, each straw represents one nesting cavity where an individual female bee could store pollen provisions and lay her eggs. Straws were placed in a 10 cm diameter PVC pipe tube, capped on one end, and covered with a corrugated plastic roof. Nests were attached at 1m height to step in fence posts and installed in a south to south-east orientation. Placement at farms was contingent on farm manager approval and typically occurred at the edges of row crops (Figure 2A), while nests were centrally installed ∼15 m into urban prairie sites (Figure 2B). Twice per month all nests were checked for nesting activity and any occupied straws were removed for further analysis and replaced with empty straws.

**Figure 2.**
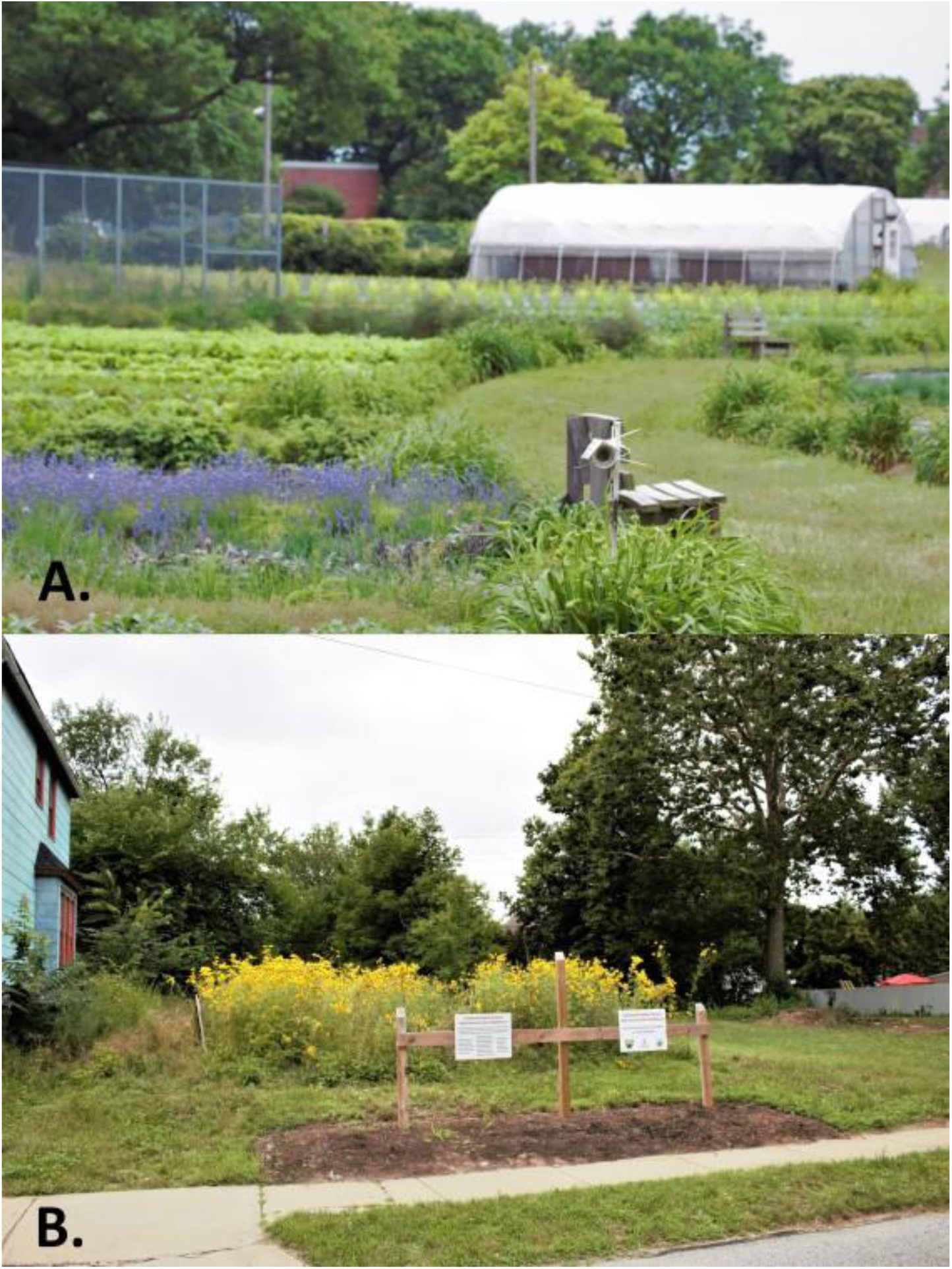
Illustrative photos of greenspace habitat types. Legacy cities, like Cleveland, Ohio, USA, contain an abundance of vacant land which can be re-purposed into unique greenspaces like (A) urban farms or (B) urban prairie conservation habitats, with implications for urban fauna.

Concurrent with each nest collection, we estimated foraging resources at all sites. We used eight 1 m^2^ quadrats positioned 5 m and 10 m from the nests. Quadrats were arranged along four perpendicular transects set at cardinal directions or, if that was not possible, arranged in an evenly spaced array across the site’s width. In each quadrat, we quantified bloom richness, and floral area. *Bloom richness* per site was estimated by summing the number of flowering species recorded from all 8 quadrats. *Bloom area* was calculated by averaging the size of each flowering species from five representative bloom size measurements and then multiplying this average bloom size by the number of floral units within a site. Floral units typically consisted of individual flowers, but complex umbellate racemes and spikes were sometimes considered as one floral unit, as described in Turo et al., 2021.

### Larvae processing and pollen library preparation

After nesting straws were collected, we x-rayed all straws for 10 seconds at 20 KV using a Faxitron LX-60 Cabinet X-ray System to confirm bee nesting and count larvae. Post X-ray analysis, straws were dissected, and pollen and larvae were separated. One larval head per nesting straw was removed for DNA extraction and Sanger sequencing at the Canadian Center of DNA Barcoding (CCDB) at the University of Guelph (Appendix 2).

Next, all pollen balls from a single nesting straw were aggregated into one pollen sample. Thus, each pollen sample represents one maternal bee’s foraging and pollen collection during nest provisioning. Pollen samples were first homogenized and then lysed in a CTAB lysis buffer using a Mini-BeadBeater 8 machine (Biospec, Bartlesville, OK). To ensure that all pollen grains were pulverized, we verified random samples under the microscope. Following pollen pulverization, we extracted DNA from a 250 ul aliquot of lysate (Appendix 3). For all samples, DNA yield and purity were verified using a NanoDrop spectrophotometer, and samples were diluted to ∼100 ng/ul.

We targeted three gene regions for sequencing: Nuclear ribosomal ITS2 and plastid *trnL* and *rbcL*. Each gene region was amplified for each sample in separate reactions using published primer sets (Appendix 3). We used a nested 3-step PCR protocol for Illumina library preparation (Appendix 3, Richardson et al., 2019; Sponsler et al., 2020) to minimize taxonomic amplification bias for each marker. Finally, all samples were normalized with a SequalPrep Normalization Plate Kit (Thermo Fisher Scientific), pooled equimolarly and sequenced using an Illumina MiSeq (2 × 300 cycles) at the Molecular and Cellular Imaging Center (MCIC) at The Ohio State University.

### Landscape composition and configuration

The Cleveland City Planning Commission provided landscape data derived from 2011 aerial imagery at a resolution of 1 m^2^. We a priori chose to characterize landscapes at 200 and 1500 m radii surrounding our sites to represent a range of foraging distances, corresponding to small and larger sized cavity nesting bees (Zurbuchen et al., 2010). A previous study in this same system found that nesting patterns for cavity nesting bees and wasps were best predicted at a scale of 1500m ( Turo & Gardiner, 2021). Landscape composition was described by percentage: 1) *impervious surface*, 2) *buildings*, 3) *grass and shrubs*, 4) *tree canopy over impervious surface*, 5) *tree canopy over buildings*, and 6) *tree canopy over grass and shrubs*. We calculated Shannon landscape diversity (*SD*) from this compositional data using FRAGSTATs v4.2 (McGarigal et al., 2012). We then re-categorized landcover data as “greenspace” (*grass and shrubs*; *tree canopy over grass and shrubs*) or “other” to quantify greenspace configuration. We used FRAGSTATs to measure the average patch isolation between greenspaces (ENN, or Euclidean Nearest Neighbor distance) and the percentage of a landscape radius that was composed by the largest patch of greenspace (LPI).

## Analysis

### Do urban bees use seeded native plants for nest provisioning?

To identify which plants urban bees foraged on while nesting, we identified bee and pollen DNA from individual nesting straws. Bee species were identified by larval DNA sequences provided by the Canadian Centre of DNA barcoding. We used the Barcode of Life Data Systems (BOLD) identification engine to compare larval sequences to species data records in BOLD. Only species previously recorded from Cleveland, OH were considered (Sivakoff et al., 2018; Turo et al., 2021; Turo & Gardiner, 2021) and identifications were considered a match if sequences were ≥ 98% similar and were assigned a species-level Barcode Index Number (BIN) from BOLD (Ratnasingham & Hebert, 2013). When a species-level match was unattainable (n=2), we identified larvae to genus (*Osmia* vs. *Megachile*) based on DNA sequences and observed nesting material, i.e., mud or leaves.

Plant species were identified after Illumina sequencing using the following bioinformatic workflow. First, paired forward and reverse reads were merged using Pear (v0.9.1; Zhang *et al*. 2014) with the base quality threshold (-q) set to 14, the minimum trim length (-t) set to 120 and a minimum assembly length (-n) of 250 bp. Given the length variability of *trnL*, only a fraction of forward and reverse reads could be assembled, so we concatenated assembled reads with unassembled forward sequences into a single file per sample for this locus. Sequences were then aligned against custom-built ITS2, *rbcL* and *trnL* reference libraries using VSEARCH semi-global alignment (v2.8.1; Rognes *et al*. 2016). Custom-built databases were curated using MetaCurator, described in Richardson *et al*. 2020, and filtered to only include species known to be present in Ohio according to the USDA Plants Database (available at https://github.com/RTRichar/MetabarcodeDBsV2). During alignment, the percent identity thresholds from Appendix 4 were used to classify sequences to family, genus or species, respectively. Only ITS2 sequences were identified to species given the poor discriminatory power of the plastid markers. Following classification using each individual marker, consensus-based median proportional abundances of each taxon were calculated for each sample at the genus and family levels using the methods provided in Richardson et al., 2019. Briefly, this analysis included an initial filtering step for the removal of any taxa present at less than 0.01 percent for any individual marker. Following this filtering, we required that taxa be detected in at least two of the three markers used and calculated the median proportional abundance of each taxon across the markers in which it was detected. Lastly, we filtered out any taxa with a median proportional abundance of less than 0.01 percent of the sample.

### Does urban bee pollen collection vary with species identity and nest location?

To test whether bees foraged on different plant communities when nesting in urban farms vs. targeted conservation greenspaces, we compared bee-collected pollen using a Permutational Multivariate Analysis of Variance (PERMANOVA) from the ‘adonis’ function in vegan (Oksanen et al., 2019; “R Development Core Team,” 2014). Pollen samples were pooled by collection period for each habitat type and visualized with the ‘metaMDS’ function (Bray-Curtis distance distribution, three dimensions).

Next, because our greenspaces varied in seeded plant diversity and because generalist bees are known to forage on diverse plants to provide nutritionally balanced pollen to their offspring (Trinkl et al., 2020; Vaudo et al., 2020), we compared pollen diversity across our nesting sites. First, we calculated Shannon diversity of pollen in individual nesting straws. Then, we used linear mixed effects models (LMM) to assess what factors drove pollen diversity patterns. Our full model included fixed effects of bee species identity, local foraging resources in the habitat (*bloom richness, bloom area*), greenspace habitat (*prairie, farm*), and landscape configuration (*SD, ENN, LPI)* at 200m and 1500m surrounding a site. Only bee species which were present in > 1 nesting straw were included in regression analyses. Also included was a term which corresponded to the Julian date within a year when a nesting straw was collected, and a random term, *neighborhood (n=6),* which accounted for spatial autocorrelation as most sites within a neighborhood were located within 1500m of each other (Appendix 1).

Prior to regression analyses, we tested for multicollinearity between variables and ensured all Variance Inflation Factors (VIF) did not exceed 3 (Zuur et al., 2009). Explanatory variables were centered and scaled. Initially, we included all possible interactions within our full model but removed these interactions as none were significant. After identifying our full model, we then conducted backwards stepwise model selection by removing the least significant term from our full model and comparing full and reduced models with a likelihood ratio test. Finally, we checked our best fit model for normality by plotting residuals vs. predicted values through package ‘DHARMa’ (Hartig, 2020). Model fit was assessed with Cox and Snell’s pseudo R^2^ through package “rcompanion” (Mangiafico, 2015).

### Is native bee fitness enhanced by targeted plantings or their configuration within the landscape?

Finally, because targeted conservation habitats are intended to support native bee populations in cities, we used generalized linear regression models to assess the nesting productivity of native bees (Russo, 2016), i.e., native larvae abundance per site, in these habitats versus urban farms. Larval data was zero-inflated and consisted of >90% zeros. Excessive zeros likely derived from bees affinity for natural nesting substrate over artificial trap nests (Everaars et al., 2011; Turo & Gardiner, 2021). To determine which model distribution (zero-inflated Poisson (ZIP) vs. zero-inflated negative binomial (ZINB)), and which scale of landscape data (200m vs. 1500m) best fit larval data we used an Akaike Information Criteria (AIC) analysis to compare all four combinations of distribution and landscape scale. Full models consisted of collection year (*2017, 2018*), a random term (*neighborhood*), and all habitat (*prairie, farm*), forage (*bloom richness, bloom area*), landscape variables (*SD, ENN, LPI*) included in previous analyses. All explanatory variables were centered and scaled to assist model convergence. The model with the lowest AICc score (±2.00) determined which model distribution and landscape scale we used. We then performed backwards model selection with likelihood ratio tests to eliminate variables from our full model. The best fit model was further evaluated for normality and dispersion with the ‘DHARMa’ package (Hartig, 2020) and we calculated model’s goodness of fit with Cox and Snell’s pseudo R^2^ through the package “rcompanion” (Mangiafico, 2015).

## Results

Over the course of two years, we collected and identified 709 larvae from 129 bee nests. Seven species of bees were identified, of which four were native and three were non-native (Table 1). One non-native bee species, *Osmia caerulescens*, comprised 76% of all larvae collected. A high abundance of *O. caerulescens* is common in urban areas such as Cleveland, OH, as this species is known to nest in masonry crevices found in urban infrastructure (Turo & Gardiner, 2021).

**Table 1.**
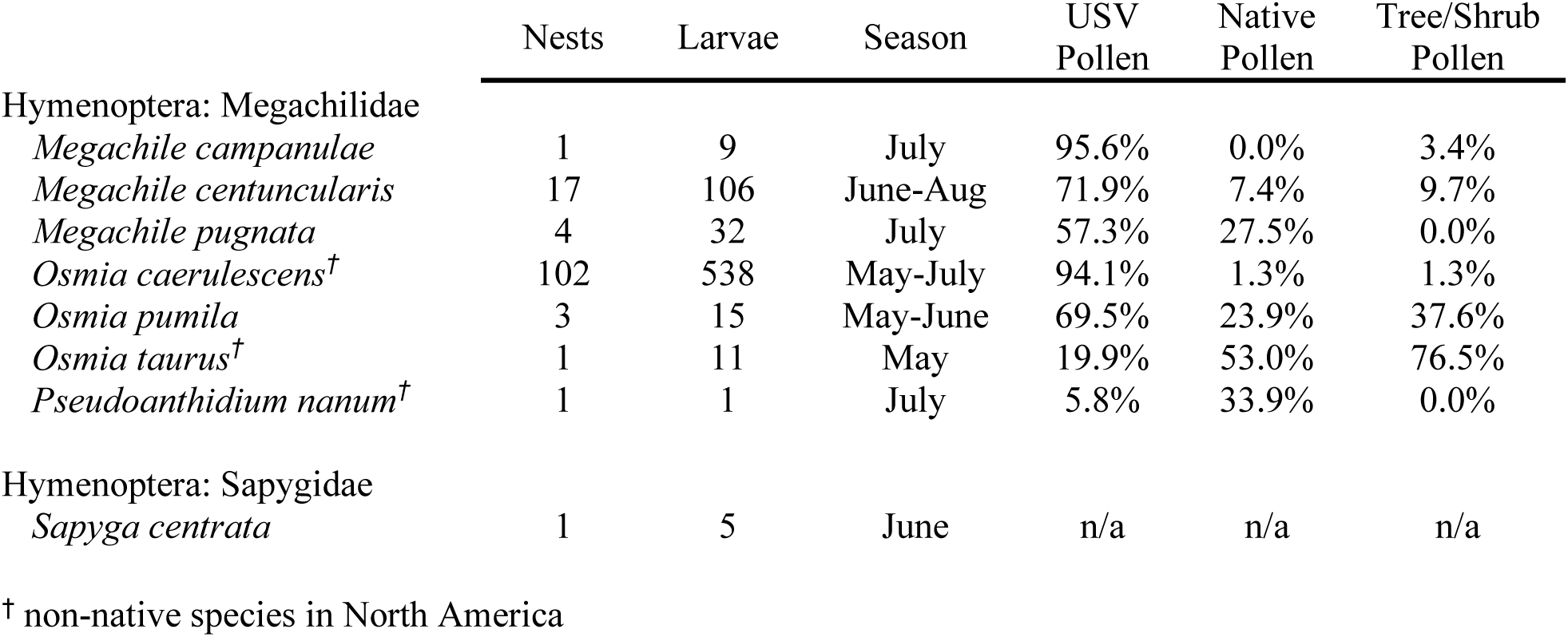
We collected and identified 709 larvae across 129 nesting straws, with most larvae collected in June and July. In addition, one parasitic wasp species, *Sapyga centrata*, was collected from an *Osmia pumila* nest. Pollen provisions varied by percent urban spontaneous vegetation (USV), native genera, and woody forage (trees/shrubs). Here, percent native pollen only pertains to a plant’s status in North America and does not indicate whether pollen originated from a native plant species within a seeded pollinator planting.

### (1) Do urban bees use seeded native plants for nest provisioning?

Metabarcoding data revealed that bees foraged on 133 plant genera, of which only 15 were abundant in quantities greater than 0.5% of all identified pollen (Appendix 5). Urban spontaneous vegetation species (USV) were observed frequently across all pollen samples (Figure 3). Roughly half (54%) of all observed pollen genera were from weedy plants while 24% of pollen genera were crops, ornamentals, or native wildflowers intentionally seeded in the urban farms or prairies. By abundance, *Trifolium* (51%), *Lotus* (11%), and *Securigera* (11%) pollens dominated *Osmia caerulescens* nesting provisions (Figure 3A) while *Lotus* (15%), *Securigera* (12%), and *Cichorium* (11%) pollens were most abundant across all native bee nesting provisions (Figure 3B-E). Conversely, we identified pollen from 9 of the 22 species of native wildflowers that were blooming in our urban prairies, but only 1% of total pollen was collected from seeded native plants (Appendix 6). Most of these foraging records were from one native bee species, *Megachile pugnata*, which frequently fed on *Ratibida* (9%), and *Rudbeckia* (6%) (Figure 3D). In general, native bees tended to collect greater proportions of seeded wildflowers, food crops, and ornamentals (Figure 3) than *O. caerulescens,* who only collected *Rubus* (1%) pollen with any frequency. Species-level patterns varied substantially, especially since *Osmia* spp. foraged in spring while *Megachile* spp. provisioned their nests in summer. For example, we note that early spring bees *Osmia pumila* (nests=3) and *Osmia taurus* (nests=1) that nested in May foraged frequently on flowering trees and shrubs (*O. pumila:* 38%, *O. taurus:* 77%) whereas pollen collected from later season bees averaged only 6% woody resources.

**Figure 3.**
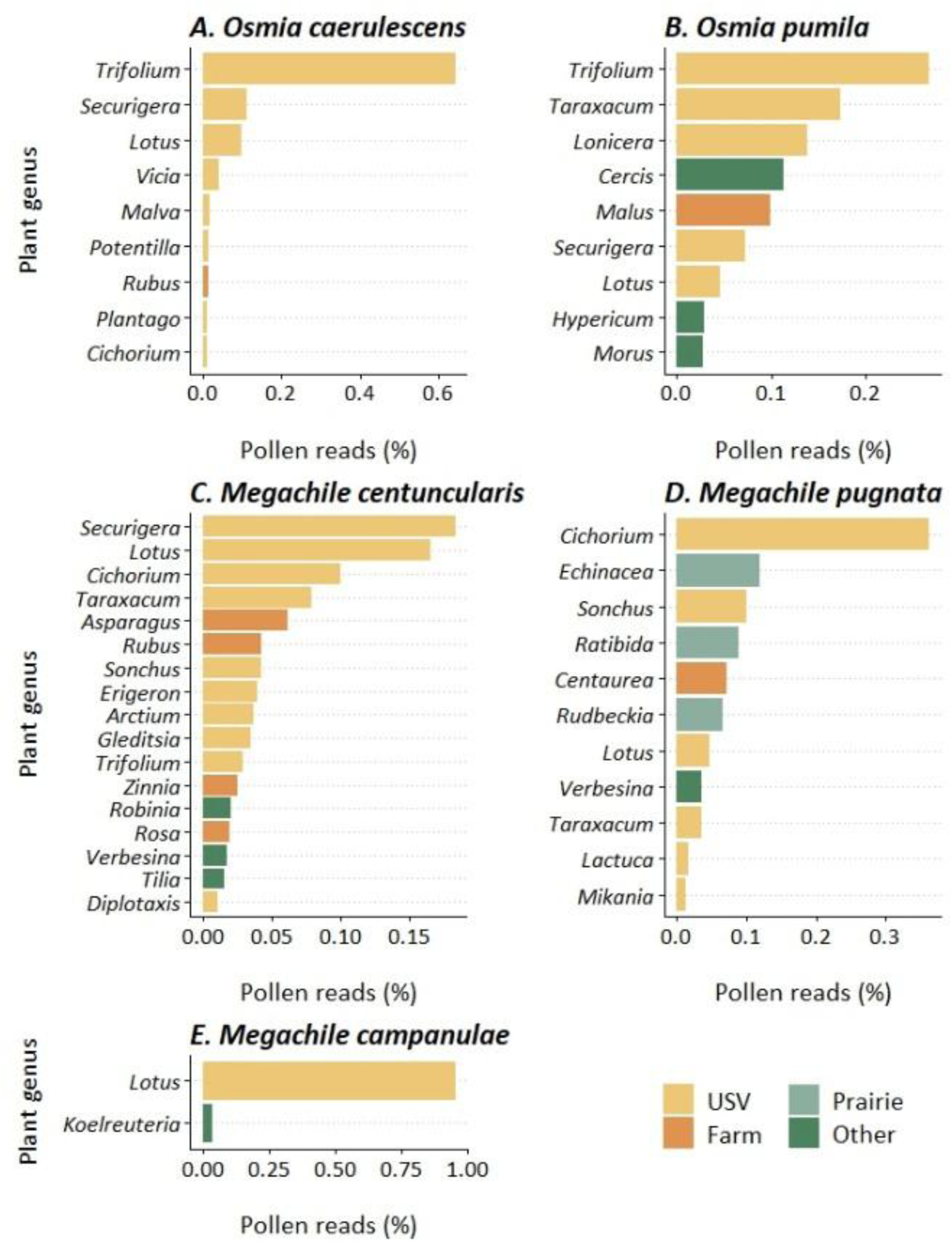
Abundance of plant genera within pollen provisions. Pollen DNA revealed dominant forage used by our most abundant bee (A) *Osmia caerulescens* (100 nests) and 25 native bee nests from 4 species: (B) *Osmia pumila* (n=3), (C) *Megachile campanulae* (n=1), (D) *Megachile centuncularis* (n=17), (E) *Megachile pugnata* (n=4). Pollen source is indicated by colors and shows whether a plant was intentionally planted in a 1) farm or 2) prairie, or whether a plant occurred spontaneously in the urban environment as a 3) weed (i.e., USV) or 4) its origin was unknown (i.e., other). Genera identifications are based on consensus-filtered, median read abundances identified from metabarcoding two plastid loci, *trnL* and *rbcL*, and one nuclear ribosomal locus, ITS2. Only plant genera contributing > 1% of pollen collected by an individual species were included.

### (2) Does urban bee pollen collection vary with species identity and nest location?

We compared bee-collected pollen from urban farms and conservation greenspaces to assess if maternal bees foraged on different plant communities when nesting in these two habitats (Figure 4). PERMANOVA analyses indicated that greenspace type did not influence the pollen composition of bee’s nesting provisions (*p* > 0.05, R^2^= 0.09).

**Figure 4.**
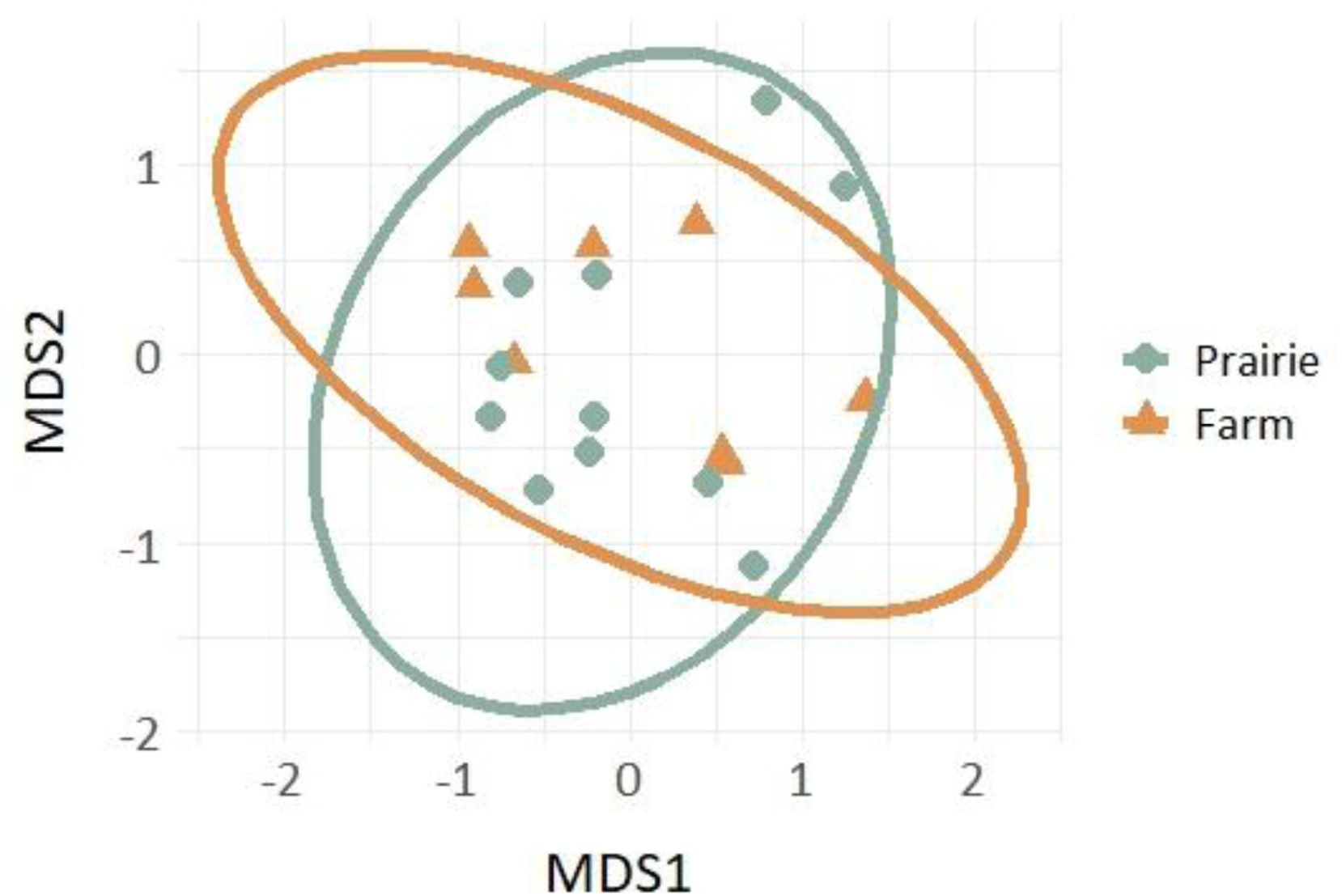
Comparison of pollen community composition for bees nesting in unique greenspace types. Nesting bees collected pollen from similar plant communities when nesting in urban farms versus targeted pollinator plantings seeded into urban prairies. Pollen data is pooled across nesting straws for each sampling period. Ellipses represent 95% confidence intervals.

Pollen diversity within a nesting straw was best explained by a bee’s taxonomic identity and by two landscape configuration metrics at 200m radii surrounding a site. Bee species was a significant predictor of pollen diversity (χ^2^ = 41.4, df = 3, *p* < 0.001), with post-hoc Tukey’s HSD tests revealing that *Osmia* spp. generally collected fewer plant genera than *Megachile* spp. (Figure 5A, Appendix 7). At a landscape scale, largest patch size was significant (χ^2^=12.0, df= 1, *p* < 0.001), and increased pollen diversity was positively correlated with a larger patch of greenspace within a 200m radius of their nest (*Coef:* 0.012 ± 0.04, Figure 5B). Likewise, greenspace patch isolation was significant (χ^2^=9.64, df= 1, *p* = 0.002) and pollen diversity was negatively correlated with patch isolation (*Coef:* −0.013 ± 0.04, Figure 5C).

**Figure 5.**
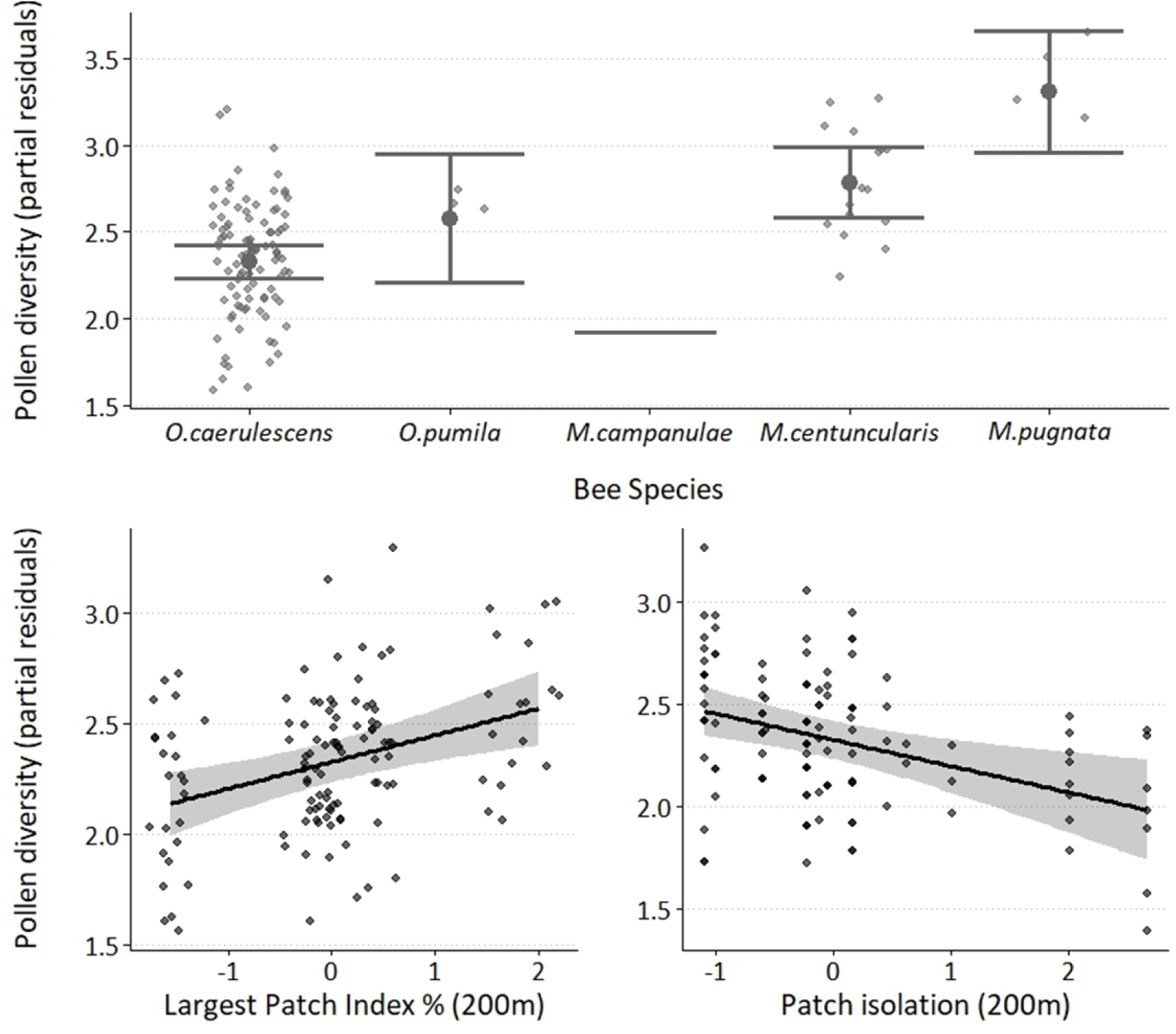
Diversity of pollen genera in a bee’s nesting provisions varied by bee species and configuration of adjacent urban greenspaces. Pollen diversity was primarily driven by (A) taxonomic identity with spring *Osmia* spp. bees tending to collect fewer plant genera than summer *Megachile* spp. (post-hoc pairwise comparisons are listed in Appendix 7). Pollen diversity also increased in (B) landscapes with a larger patch of greenspace within 200m of a bee’s nest, and in (C) landscapes with decreased patch isolation within 200m of a nest. *Megachile campanulae*, *Osmia taurus*, and *Pseudoanthidium nanum* were excluded from regression analysis due to their low abundance (n=1). However, *M. campanulae* was included in visualization to show variance in native bee’s pollen collection. Partial residuals indicate contribution of individual variables towards model variance.

All other included variables were not significant and were subsequently removed during backwards model selection. We assessed our best model’s goodness-of-fit with a Cox and Snell’s pseudo R^2^ (_pseudo_ R^2^ = 0.33). We also validated our best fit model by comparing it to an intercept only model with a likelihood ratio test (χ^2^=49.82, df= 5, *p* < 0.001).

### (3) Is native bee fitness enhanced by targeted plantings or their configuration within the landscape?

Native larvae abundance per site was best fit with a zero-inflated Poisson (ZIP) distribution with landscape features at 1500m scale, according to AICc analysis. Our best fit ZIP model consisted of a binomial model, which described patterns contributing to presence of larvae within a nest, linked to a count model with a Poisson distribution, which described native larvae abundance, if present. We found that the presence of native larvae at a site was negatively associated with bloom richness surrounding a nesting site (*z* = −3.36, *p* <0.001, Table 2). When present, native larvae abundance decreased with bloom richness (*z* = −7.39, *p* < 0.001, Figure 6A), and increased with greater bloom area (*z* = 2.59, *p* = 0.010, Figure 6B). Larvae were more abundant in urban farms vs. urban prairies (*z* = 7.43, *p* < 0.001, Figure 6C) and in 2017 collections versus 2018 collections (*z* = −5.58, *p* < 0.001, Figure 6D). Finally, landscape configuration at 1500m influenced native larvae abundance with more larvae occurring in landscapes with less isolation between greenspace patches (*z* = −3.64, *p* < 0.001, Figure 6E). We also compared our best fit ZIP model to an intercept only model with a likelihood ratio test, which indicated a strongly significant improvement of fit (χ^2^=107.07, df= 8, *p* < 0.001).

**Figure 6.**
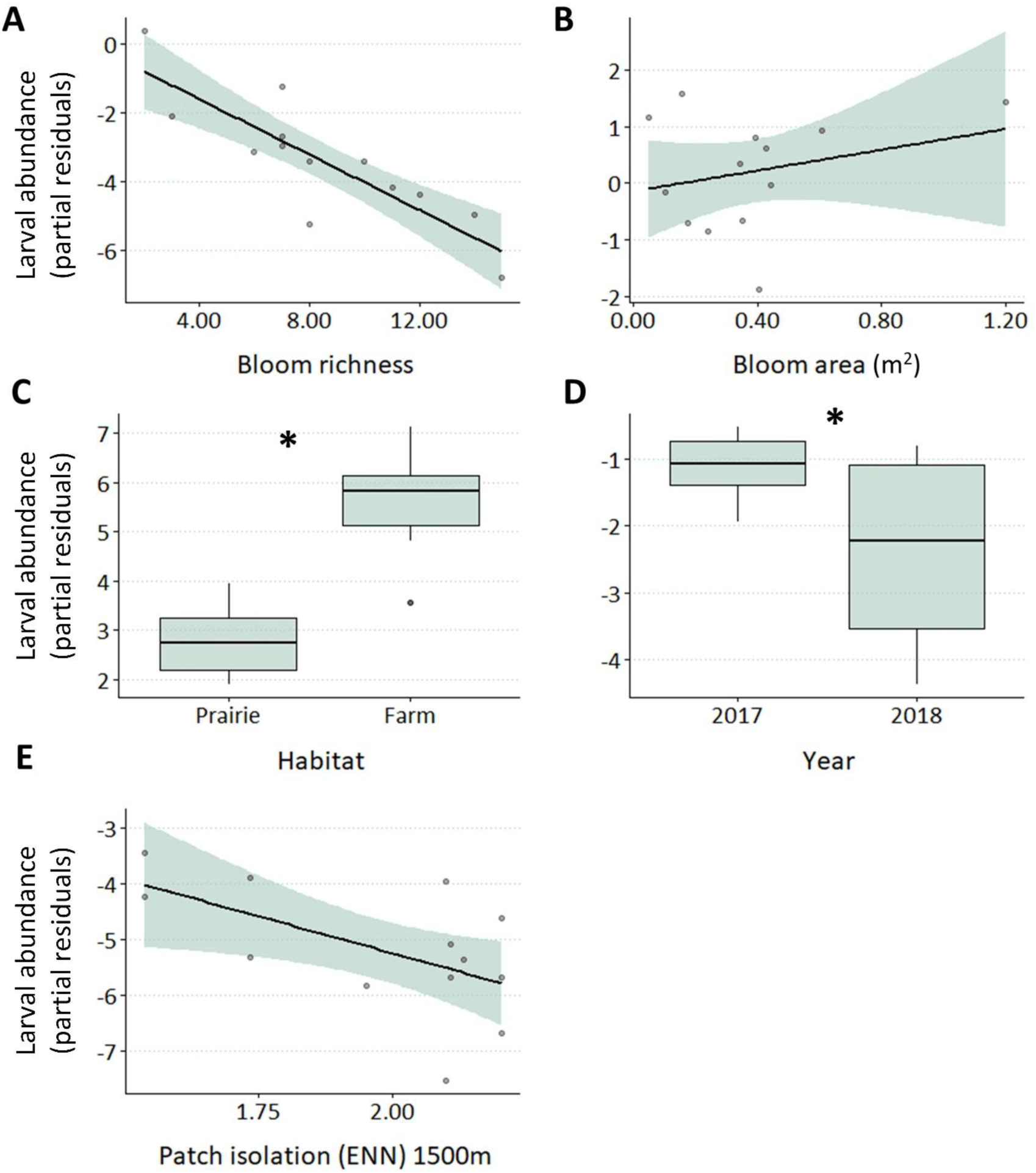
Partial residual plots of best fit ZIP model for native bee larvae abundances in urban farms and pocket prairies. Native larvae abundance was correlated with increased bloom area but decreased bloom richness. Native larvae were also more abundant on farms and in 2017. As greenspace patches became more isolated (ENN) in a 1500 m radius surrounding a nesting site, native larvae were less abundant. Partial residuals indicate contribution of individual variables towards model variance. Shaded bands indicate 95% confidence intervals.

**Table 2.** Results from a composite, Zero-inflated Poisson (ZIP) model which collectively describes the presence of native larvae in a trap (binomial model), and then the abundance of native larvae, when nesting occurred (count model). ZIP model goodness-of-fit was evaluated with Cox and Snell’s pseudo R^2^ (0.31%). Parameters for the binomial component are reported on the log-odds (i.e., logit) scale, while parameters for the Poisson component are reported on the log scale.

| ZIP model variables | Estimate | SE | *z* | 95% CI | *p* |
| --- | --- | --- | --- | --- | --- |
| Presence in trap nests (binomial) |  |  |  |  |  |
| *intercept* | 3.65 | 0.51 | 7.15 | 2.65, 4.65 | < 0.001* |
| *bloom richness* | -1.15 | 0.34 | -3.36 | -1.82, -0.48 | < 0.001* |
| Abundance in trap nests (count) |  |  |  |  |  |
| *intercept* | 0.67 | 0.57 | 1.18 | -0.45, 1.79 | 0.239 |
| *habitat (prairie, farm)* | 2.88 | 0.39 | 7.43 | 2.12, 3.64 | < 0.001* |
| *year (2017,2018)* | -1.19 | 0.21 | -5.58 | -1.61, -0.77 | < 0.001* |
| *bloom area* | 0.37 | 0.14 | 2.59 | 0.09, 0.65 | 0.010* |
| *bloom richness* | -1.27 | 0.17 | -7.39 | -1.60, -0.93 | < 0.001* |
| *patch isolation (ENN)* | -0.67 | 0.18 | -3.64 | -1.03, -0.31 | < 0.001* |

bloom richness= average bloom richness within a greenspace patch; habitat = greenspace type where nest was located (urban prairie vs. urban farm); year= year of collection (2017 vs. 2018); bloom area= average bloom area (mm<sup>2</sup>) within a greenspace patch; patch isolation (ENN)= average distance (m) between greenspace patches in a landscape (1500m radius)

## Discussion

Here, we evaluated how bee nest provisioning and reproduction varied with greenspace type (targeted conservation habitat vs. multi-service urban farms) and surrounding landscape structure. Pollen DNA documented that urban cavity nesting bees predominately used urban spontaneous vegetation (USV) as forage, especially pollen from the Fabaceae family. USV pollens were abundant in most nesting straws regardless of bee identity, landscape context, or nesting site location within an urban farm or native prairie planting, suggesting that urban weeds are broadly beneficial to city bees. Likewise, our targeted conservation habitats were less effective than multi-service urban farms in supporting native cavity nesting bee reproduction. This suggests that multi-service greenspaces (e.g., urban farms) may promote win-win conservation scenarios for people and pollinators. Lastly, we confirm that any greenspace’s value to bees also relies, in part, on landscape configuration as greater patch connectivity allows access to diverse forage and can improve nesting success.

We established our urban prairies with the intention of providing bee forage, yet DNA revealed that cavity nesting bees generally foraged on adjacent weeds and not on the flowering plants we seeded. In many ways, this was unsurprising as similar weedy foraging has been observed in other studies, especially to *Trifolium* (clover) (Larson et al., 2014; Lerman et al., 2018; MacIvor et al., 2014; Turo et al., 2021— but see Casanelles-Abella et al., 2022). Likewise, we expected frequent foraging on urban spontaneous vegetation because several weeds in the Fabaceae family frequently grow in Cleveland (Turo et al. 2021) and most of the bees we collected were *Osmia* spp., which commonly collect pollen from Fabaceae plants (MacIvor et al., 2014; Voulgari-Kokota et al., 2019) that exhibit high protein to lipid ratios (Vaudo et al., 2020). Weedy diets were not exclusive to *Osmia* spp. or even our most abundant exotic bee, *Osmia caerulescens* though; each bee taxa that we recruited used a weedy plant species (e.g., *Trifolium, Securigera, Lotus, Cichorium)* as their dominant pollen source (Figure 3). This suggests to us that the ability to forage on abundant weedy plants might serve as a biotic filter (Aronson et al. 2016) for which bee species persist in urban environments. Moreover, if a greenspace’s goal is to support pollinators which are already present in cities and providing pollination services, land managers should consider adopting a socially-responsible (Turo & Gardiner, 2020), infrequent mowing schedule that allows urban spontaneous vegetation to bloom and provide forage for urban bees (Lerman et al., 2018).

Still, we did not expect cavity nesting bees would exhibit such limited interest in our seeded native plants (Figure 3, Appendix 6, 7), as observational studies in this same urban prairie system, recorded much higher visitation rates (6-8%) of bees on seeded native plants (Turo et al. 2021) than we document here with pollen DNA. For this study, despite 2 additional years of prairie plant establishment and floral growth, seeded native plants comprised < 1% of all collected pollen, seeded native plants were only a dominant pollen source for one native bee species, *Megachile pugnata* (Figure 3) Phenology accounts for some of this discrepancy. We collected 46/129 nests before June 15^th^, but most of the seeded plants, except *Coreopsis tinctoria*, tended to bloom later in the season and were not available to early season bees. In addition, we only collected pollen from cavity-nesting bees, which represents a subset of the bee community, and the potential benefit of seeded plants to this guild may be less than for other subsets of the bee community. For example, in another observational study in this prairie network, we observed bumble bees as frequent foragers on seeded native plants, especially in late summer, with 78% of *Bombus* spp. foraging records in pollinator plantings occurring on seeded native plants (Pham et al. *in revision*). Thus, we do not conclude that the prairies were wholly ineffective, but rather their benefit as forage is contingent on time since establishment and is likely targeted at different taxa than those represented in this study. Finally, we note that our molecular analysis showed that when cavity nesting bees did feed on native plants, trees were common pollen sources (Figure 3, Table 1), representing 8/15 of the top native plant genera foraged on (Appendix 8). This highlights the importance of pollen DNA in revealing difficult-to-observe foraging (Bell et al., 2023), and aligns our study with three recent metabarcoding studies, which document urban bees feeding on a mixture of weeds, ornamental forbs, and native trees (Casanelles-Abella et al., 2022; Dürrbaum et al., 2023; Potter et al., 2019). In addition, frequent urban tree foraging indicates that any greenspace intending to support urban pollinators, whether for conservation or ecosystem service provision, should consider diversifying new plant installations to include woody species.

Our second key finding was that native cavity-nesting bees’ reproductive success was greatest when nesting at urban farms. This is promising because it suggests that multi-service urban greenspaces still benefit native bee biodiversity, even if their primary goal was not pollinator conservation. From an urban planning perspective, this is particularly significant because social and financial support can be more available for multi-service greenspaces than targeted conservation spaces. As continued social and financial investment is critical for long-term management of an urban greenspace (Turo and Gardiner 2019, 2020), it is possible that urban farms might be better suited to achieve conservation goals in some cities. In part, urban farms may perform better long-term because pollinator plantings are frequently abandoned (Turo & Gardiner, 2019), especially if funded by short-term grants or if site management is not explicitly budgeted for during project onset. Conversely, urban farms can have more stable funding mechanisms by generating their own income, and there are more urban farms already present in cities worldwide, as 800 million farmers currently grow crops in urban environments (FAO, UN).

Why might urban farms achieve better outcomes for cavity nesting bees though? In our study, there are several ecological reasons potentially driving greater native larvae abundances in urban farms. First, we were unable to control the size of available urban farms so that prairies and farms represented equivalent habitat area. It is possible that the urban farms attracted more nesting because these greenspaces were larger. Certainly, patch size is a consistent positive driver of bee abundance, bee richness, and cavity-nesting bee abundance in cities and suburban areas in Europe, Asia, and North America (Banaszak-Cibicka et al., 2016; Pardee & Philpott, 2014; Quistberg et al., 2016; Stewart et al., 2018). However, in our analyses we did not detect an effect of greenspace patch size at 200m radius on larvae abundances—complicating this interpretation. It is also possible that the urban farms offered more flowering resources to bees, or different types of flowering resources that contributed to greater nesting success. Floral abundance is frequently cited as a main driver of bee abundances in urban environments (Hyjazie & Sargent, 2022; Winfree et al., 2011). But, in our study, DNA revealed that pollen provisions were not compositionally different in farms versus pollinator plantings (Figure 4) and urban farms were weakly correlated with decreased bloom area (r= −0.010). Our best interpretation, therefore, is that urban farms may have hosted pre-existing populations of cavity nesting bees, possibly because they are more established habitats or because they contain greater nesting resources (Felderhoff et al., 2023), like raspberry canes. While we did not quantity nesting habitat in our study, anecdotally, the urban farms frequently grew raspberry plants and were less disturbed. For example, all prairies were mown once a year in the Fall, and this management might have reduced nesting resource availability.

Finally, landscape configuration also influenced urban bee nesting. Greater patch connectivity correlated with increased native bee larvae and higher pollen diversity in nests. These results align with previous research and indicate that larger, more connected patches of greenspace may allow for greater access to diverse foraging resources available in the urban environment. For example, greater patch connectivity benefits resource capture by bees (Van Rossum & Triest, 2012), while urban fragmentation is known to limit bee dispersal and foraging decisions (Bhattacharya et al., 2003; Harrison & Winfree, 2015; O’Connell et al., 2021). Likewise, cavity nesting bees are documented as incorporating greater pollen richness in their diets as nests are placed further from city centers, where fragmentation and impervious surface are greater (Dürrbaum et al., 2023). Of note, different scales of landscape structure influenced nesting success (1500 m) and bee’s foraging (200 m), suggesting that urban bees may disperse longer distances when locating an optimum nesting site, but then provision these nests via foraging at smaller scales. Similar patterns have been identified in other urban environments, where bumble bees foraged on much smaller scales than they are capable of dispersing (O’Connell et al., 2021). Such scale differences also support the conjecture that nesting resources may be more limiting than foraging resources within cities (Sivakoff et al., 2018; Stewart et al., 2018). The implication of this is that while most pollinator conservation actions focus on creating more forage, bees would likely benefit most from increased nesting habitat (Felderhoff et al., 2023)—although this a difficult task to undertake as there is even less guidance about how to design a habitat for bee nesting than for bee foraging.

## Conclusions

Pollen DNA confirmed several observational datasets that urban spontaneous vegetation widely supports urban bee foraging (Larson et al., 2014; Lowenstein et al., 2019; Turo et al., 2021), suggesting that greenspace management practices which tolerate weeds are beneficial to city-dwelling bees. We did not find that targeted conservation plantings conferred a fitness benefit to nesting native bees or were frequently fed on by most bees. Rather, multi-service urban farms harbored more native bee larvae and achieve the complimentary goals of providing pollinator habitat and urban food production. However, we emphasize that the value of a pollinator planting depends on which taxa it aims to support, and targeted conservation plantings may still benefit different guilds of bees not considered in this study. Finally, the landscape scale patterns in our study suggest that urban fragmentation structures plant-pollinator interactions. Maintaining a connected network of greenspace, and maximizing patch sizes, is a critical consideration when creating urban pollinator habitat (see also Banaszak-Cibicka et al. 2016, Turo & Gardiner, 2021). As such, situating new urban pollinator habitats adjacent to pre-existing, larger greenspaces and improved habitat connectivity (Braaker et al., 2014; Van Rossum & Triest, 2012) are important considerations for future urban greening.

## Notes

### Competing Interest Statement

The authors have declared no competing interest.

