## Supplementary Material for "Pollen DNA reveals urban cavity-nesting bees feed primarily on weeds and frequently nest in urban farms"

### Appendix 1. Greenspace design and management

High diversity (22 forbs, 3 grasses) and medium diversity (12 forbs, 3 grasses) urban pocket prairies were seeded within 30 x 12 m vacant lots of Cleveland, Ohio in 2014 by Ohio Prairie Nursery (Hiram, OH). Vacant lots were leased by The Ohio State University on behalf of Dr. Mary Gardiner from the Cleveland City Land Bank. Prior to prairie establishment, all vacant lots were turf grass habitats, following demolition of an abandoned residential property. Our two prairie treatment designs were established in six neighborhoods (Buckeye, Central, Detroit Shoreway, Fairfax, Hough, Slavic Village) in a randomized block design. Throughout this study, all prairies were mown once per year in the Fall and site edges were maintained with monthly mowing and trimming. See Turo et al. 2021 for more detail on pocket prairie establishment. Finally, although high and medium diversity prairies were initially intended as unique greenspace designs, plant establishment was poor over the course of the study (see Appendix 6), and most seeded species did not bloom or were rare. In effect, all prairies, regardless of seeded diversity contained a similar set of blooming flowers and were thus treated as the same type of greenspace in subsequent analysis. The only exception to this general rule was *Coreopsis lanceolata* which bloomed only in half of the prairie sites, although not at high abundances.

Urban farms were also developed on previously vacant land and were situated in the same six neighborhoods as pocket prairie conservation sites. Farms were included on a case-by-case basis depending on availability, and, whenever possible, two farms were included in each neighborhood (n=2: Buckeye, Detroit Shoreway, Hough; n=1: Central, Fairfax, Slavic Village). Farms were managed by Bay Branch Farm, Cleveland Botanical Gardens, Cuyahoga County Extension, Kinsman Farm, and Ohio City Farm. All farms focused on vegetable production and varied in size from 2.43 ha to 0.10 ha. Ornamental flowers were frequently planted at farms in adjacent raised beds or in the field edges but flowering species

| **Medium diversity urban prairie** | **High diversity urban prairie** |
| --- | --- |
| Prairie grasses | Prairie grasses |
| *Elymus canadensis* | *Elymus canadensis* |
| *Schizachyrium scoparium* | *Schizachyrium scoparium* |
| *Sorghastrum nutans* | *Sorghastrum nutans* |
| Prairie forbs | Prairie forbs |
| *Aster novae-angliae* | *Aster novae-angliae* |
| *Bidens aristosa*[†] | *Bidens aristosa*[†] |
| *Chamaecrista fasciculata*[†] | *Chamaecrista fasciculata*[†] |
| *Eupatorium purpureum* | *Coreopisis lanceolata* |
| *Liatris spicata* | *Coreopsis tinctorial*[†] |
| *Lobelia siphilitica* | *Eryngium yuccifolium* |
| *Monarda citriodora*[†] | *Eupatorium purpureum* |
| *Monarda fistulosa* | *Gaillardia pulchella*[†] |
| *Ratibida pinnata* | *Liatris spicata* |
| *Rudbeckia hirta*[†] | *Lobelia siphilitica* |
| *Silphium perfoliatum* | *Monarda citriodora*[†] |
| *Zizia aurea* | *Monarda fistulosa* |
| | *Penstemon digitalis* |
| | *Ratibida pinnata* |
| | *Rudbeckia hirta*[†] |
| | *Silphium perfoliatum* |
| | *Silphium terebinthinaceum* |
| | *Solidago riddellii* |
| | *Tradescantia ohiensis* |
| | *Verbena hastata* |
| | *Vernonia fasciculata* |
| | *Zizia aurea* |

[†] species added as an overseed mixture in 2016 during prairie's establishment phase

**Appendix 2. Primer sets for pollen and bee metabarcoding.**

Three markers (*rbcL*, *trnL*, ITS2) were used to amplify pollen DNA from plastid and nuclear ribosomal loci. Published primer sets were obtained from (Chen et al., 2010; Kress & Erickson, 2007; Palmieri et al., 2009; Taberlet et al., 1991; White et al., 1990), and additional methodological information is available in the supplementary materials of Richardson et al. 2021. Bee metabarcoding and identification was completed at the Canadian Centre for DNA barcoding where a 658 bp gene region of the Cytochrome c oxidase subunit I (COI) was targeted using a universal insect primer set.

| Marker | Primer Sequence | Reference |
|---|---|---|
| *rbcL* | | |
| rbcL2 | TGGCAGCATTYCGAGTAACTC | Kress and Erickson 2007, Palmieri et al. 2009 |
| rbcLa-R | GTAAAATCAAGTCCACCRCG | |
| *trnL* | | |
| A49325 | CGAAATCGGTAGACGCTACG | Taberlet et al. 1991 |
| B49863 | GGGGATAGAGGGACTTGAAC | |
| ITS2 | | |
| ITS-S2F | ATGCGATACTTGGTGTGAAT | White et al. 1990, Chen et al. 2010 |
| ITS4R | TCCTCCGCTTATTGATATGC | |
| CO1 | | |
| C_LepFolF | RKTCAACMAATCATAAAGATATTGG | Canadian Centre for DNA Barcoding |
| C_LepFolR | TAAACTTCWGGRTGWCCAAAAAATCA | |

**Appendix 3. Molecular workflow including PCR conditions and additives for each marker and PCR step**

*Pollen sample preparation*

Pooled pollen samples from an individual nesting straw were weighed and a 1:10 parts solution of pollen to 70% ethanol was created. We then added three steel beads (4 mm) to pollen sample tubes and vortexed until samples were homogenized. Those samples with large pellets of pollen which did not break up during vortexing were warmed on a heat block at 65 °C for 10 minutes to facilitate homogenization. After homogenization, a 750 ul aliquot was drawn from each sample, placed in a 2 mL bead mill tube (BioSpec, Bartlesville, OK) and centrifuged at 10,000 rcf for 3 minutes. Supernatant was discarded and tubes were left for 30 minutes in the fume hood to further evaporate ethanol. Following evaporation, we added 1.25 ml of CTAB lysis buffer (100 mM Tris–HCL, pH 8.0; 2% CTAB; 20 mM EDTA) and ~0.5 ml of zirconia beads (0.7 mm Biospec, Bartlesville, OK) to each tube. Pollen samples were then pulverized for 5 minutes on high utilizing a Mini-BeadBeater 8 machine (Biospec, Bartlesville, OK) to lyse all pollen grains. Importantly, prior to DNA extraction, random pulverized samples were verified under the microscope to ensure that all pollen grains were appropriately lysed.

*Pollen library preparation and sequencing*

Following pollen pulverization, we extracted DNA from a 250 ul aliquot of lysate. Lysate was vortexed for 40 seconds in a 1.5 ml microcentrifuge tube containing 100 ul phenol, 100 ul chloroform, and 6% SiO2 phase lock grease. Organic and aqueous layers were then separated by centrifuging tubes at 16,000 rcf for 20 minutes and 200uL of the resulting aqueous layer was transferred to a new 1.5 mL tube. DNA was precipitated by adding 1 uL glycogen, 100 uL of 5M $NH_4OAc$ and 750 uL 100% EtOH, inverting the tubes, and incubating them at -20° C overnight. Tubes were then centrifuged for 20 minutes at 16,000 rcf and resultant DNA pellets were washed twice with 125 uL of 70% EtOH. Following this, 75 uL of water at 65° C was added to dissolve pellets for future storage and analysis. Pellets which did not immediately dissolve were inverted and placed on a heat block at 65° C for ~10 minutes. For all samples, DNA yield and purity were verified using a NanoDrop spectrophotometer, and samples were diluted to ~100 ng/ul.

We amplified DNA for three gene regions, nuclear ribosomal ITS2 and plastid *trnL* and *rbcL*, for each sample in separate reactions using published primer sets (Chen et al., 2010; Kress & Erickson, 2007; Palmieri, Bozza, & Giongo, 2009; Taberlet, Gielly, Pautou, & Bouvet, 1991; White, Bruns, Lee, & Taylor, 1990, Appendix 2). We employed a nested 3-step PCR protocol for Illumina library preparation (Richardson et al., 2019; Sponsler et al., 2020) to minimize taxonomic amplification bias for each marker. Our multi-locus approach enabled consensus-based filtering of DNA records and more accurate quantification of pollen DNA (Richardson et al., 2019, 2021)**.** PCR conditions and additives are listed below for each gene region and PCR step. In step one, we amplified our gene region of interest with target primers. In step two, Illumina read priming sequences were appended to DNA fragments. Finally, in step three, we appended Illumina hybridization oligos with unique dual index sample codes. Finally, all samples were normalized with a SequalPrep Normalization Plate Kit (Thermo Fisher Scientific), pooled equimolarly and sequenced using an Illumina MiSeq (2 × 300 cycles) at the Molecular and Cellular Imaging Center (MCIC) at The Ohio State University.

**Table 1. PCR conditions and additives.**

| Marker | Reaction Step | Primer conc. | Stage 1 (°C\|sec) | Stage 2 (°C\|sec) | Annealing Stage (°C\|sec) | Stage 4 (°C\|sec) | # Cycles | Stage 5 (°C\|sec) | Additives |
|---|---|---|---|---|---|---|---|---|---|
| *rbcL* | 1 | 0.5 uM | 98\|30 | 98\|10 | 58.0\|10 | 72\|15 | 20 | 72\|300 | 0.8 ul BSA, 1 ul DMSO |
| *rbcL* | 2 | 0.1 uM | 98\|30 | 98\|10 | 50.2\|10 | 72\|15 | 15 | 72\|300 | 0.8 ul BSA, 1 ul DMSO |
| *rbcL* | 3 | 0.5 uM | 98\|30 | 98\|10 | 66\|10 | 72\|15 | 25 | 72\|300 | - |
| *trnL* | 1 | 0.5 uM | 98\|30 | 98\|10 | 58.0\|10 | 72\|15 | 25 | 72\|300 | 0.8 ul BSA, 1 ul DMSO |
| *trnL* | 2 | 0.1 uM | 98\|30 | 98\|10 | 55.4\|10 | 72\|15 | 18 | 72\|300 | 0.8 ul BSA, 1 ul DMSO |
| *trnL* | 3 | 0.5 uM | 98\|30 | 98\|10 | 66\|10 | 72\|15 | 25 | 72\|300 | - |
| ITS2 | 1 | 0.5 uM | 98\|30 | 98\|10 | 60.0\|10 | 72\|15 | 20 | 72\|300 | 0.8 ul BSA, 1 ul DMSO |
| ITS2 | 2 | 0.1 uM | 98\|30 | 98\|10 | 52.3\|10 | 72\|15 | 15 | 72\|300 | 0.8 ul BSA, 1 ul DMSO |
| ITS2 | 3 | 0.5 uM | 98\|30 | 98\|10 | 66\|10 | 72\|15 | 25 | 72\|300 | - |

BSA= bovine serum albumin; DMSO= dimethyl sulfoxid

**Appendix 4**. **Alignment percent identity thresholds by marker and taxonomic rank.**

Pollen sequencing resulted in an average and standard error of 84,700 ± 2,400 plant sequences per sample across all three markers. There was significant variation in average sequencing coverage across markers ($p < 0.001$, OLS Regression), with ITS2 (39,564 ± 1,379 plant reads) yielding significantly more coverage than *rbcL* (30,914 ± 1,950) or *trnL* (14,226 ± 1,950).

| | Family | Genus | Species |
|---|---|---|---|
| ITS2 | 90 | 95 | 98 |
| *rbcL* | 95 | 97 | NA |
| *trnL* | 95 | 97 | NA |

### Appendix 5. Plant genera identified from pollen provisions in bee's nests

Proportion of identified DNA and frequency of occurrence within 133 bee's nests are noted for each plant genus. Typical growing habit (*herb, forb, vine, shrub, tree)* and common names are also listed. Common names indicate a representative species for each genera and are based on recorded plant species lists in Cleveland (Turo et al., 2021; Turo & Gardiner, 2021) and observed bloom data. Origin reflects whether a plant species corresponding to an observed pollen genus, was intentionally seeded and observed in our study habitats (*urban farm, urban prairie*), was common, urban spontaneous vegetation (Turo et al. 2021), or was not observed in our study system and grew elsewhere (residential gardens, parks, etc.). Only plants which were present in greater than 0.2% of total identified pollen are shown. † indicates plants from urban prairie seed treatments.

| Genus | Reads (%) | Count | Origin | Habit | Common name |
|---|---|---|---|---|---|
| *Trifolium* | 51.3% | 133 | USV | herb | clover |
| *Lotus* | 11.0% | 124 | USV | herb | birds foot trefoil |
| *Securigera* | 11.0% | 124 | USV | forb | crown vetch |
| *Cichorium* | 3.3% | 84 | USV | forb | chicory |
| *Vicia* | 3.1% | 55 | USV | forb | hairy vetch |
| *Taraxacum* | 2.5% | 66 | USV | herb | dandelion |
| *Rubus* | 1.4% | 26 | urban farm | shrub | raspberry |
| *Malva* | 1.2% | 67 | USV | forb | mallow |
| *Potentilla* | 1.0% | 46 | USV | forb | cinquefoil |
| *Zinnia* | 0.9% | 4 | urban farm | forb | zinnia |
| *Sonchus* | 0.9% | 38 | USV | forb | sow thistle |
| *Plantago* | 0.9% | 103 | USV | herb | narrow leaf plantain |
| *Asparagus* | 0.8% | 10 | urban farm | herb | asparagus |
| *Pinus* | 0.7% | 37 | other | tree | pine |
| *Erigeron* | 0.7% | 33 | USV | forb | daisy fleabane |
| *Papaver* | 0.5% | 4 | urban farm | forb | poppy |
| *Gleditsia* | 0.5% | 9 | other | tree | honey locust |
| *Arctium* | 0.5% | 13 | USV | forb | burdock |
| *Verbesina* | 0.5% | 7 | other | forb | wingstem |
| *Echinacea* | 0.5% | 5 | urban prairie | forb | purple coneflower† |
| *Ratibida* | 0.4% | 21 | urban prairie | forb | grey headed coneflower† |
| *Quercus* | 0.4% | 12 | other | tree | oak |
| *Helianthus* | 0.4% | 3 | urban farm | forb | sunflower |
| *Parthenocissus* | 0.3% | 5 | USV | vine | Virginia creeper |
| *Nepeta* | 0.3% | 9 | other | forb | catmint |
| *Lonicera* | 0.3% | 4 | other | shrub | honeysuckle |
| *Ligustrum* | 0.3% | 16 | USV | shrub | privet |
| *Cercis* | 0.3% | 4 | other | tree | redbud |
| *Robinia* | 0.2% | 3 | other | tree | black locust |
| *Malus* | 0.2% | 5 | urban farm | tree | apple |

| | | | | | |
|---|---|---|---|---|---|
| *Rosa* | 0.2% | 12 | urban farm | shrub | rose |
| *Centaurea* | 0.2% | 14 | urban farm | forb | cornflower |
| *Hypericum* | 0.2% | 28 | other | forb | St. John’s wort |
| *Lamium* | 0.2% | 4 | USV | herb | dead nettle |
| *Tilia* | 0.2% | 22 | other | tree | linden |
| *Rudbeckia* | 0.2% | 12 | urban prairie | forb | black eyed Susan† |
| *Chaenomeles* | 0.2% | 2 | other | shrub | flowering quince |
| *Penstemon* | 0.2% | 15 | urban prairie | forb | foxglove beardtongue† |
| *Symphyotrichum* | 0.2% | 13 | urban prairie | forb | New England aster† |

† intentionally seeded wildflower plant

**Appendix 6. Seeded plant establishment and bee foraging.**

Twenty-two species of native prairie forbs were seeded in vacant lot greenspaces. After four years of establishment, only 13 species bloomed, and few plants were flowering in appreciable amounts. Percent bloom area indicates the percentage of all sampled bloom area from May through July that each prairie species composed. Grey headed coneflower, *Ratibida pinnata*, growth exceeded all other planted species and comprised 30% of all sampled bloom area. However, percentage pollen which bees collected from plant genera corresponding to our seeded prairie plants was low.

| | bloomed? | % bloom area | foraged on? | % pollen |
|---|---|---|---|---|
| *Aster novae-angliae* | yes | < 0.01% | yes | 0.152% |
| *Bidens aristosa*† | no | - | - | - |
| *Chamaecrista fasciculata*† | yes | < 0.01% | yes | 0.000% |
| *Coreopisis lanceolata* | yes | 3.77% | yes | 0.043% |
| *Coreopsis tinctorial*† | no | - | - | - |
| *Eryngium yuccifolium* | yes | < 0.01% | no | - |
| *Eupatorium purpureum* | no | - | yes | 0.004% |
| *Gaillardia pulchella*† | no | - | - | - |
| *Liatris spicata* | no | - | yes | 0.005% |
| *Lobelia siphilitica* | no | - | - | - |
| *Monarda citriodora*† | no | 0.01% | no | - |
| *Monarda fistulosa* | yes | 5.38% | yes | 0.004% |
| *Penstemon digitalis* | yes | 0.16% | yes | 0.155% |
| *Ratibida pinnata* | yes | 30.57% | yes | 0.425% |
| *Rudbeckia hirta*† | yes | 1.10% | yes | 0.206% |
| *Silphium perfoliatum* | yes | 0.89% | yes | 0.007% |
| *Silphium terebinthinaceum* | no | - | - | - |
| *Solidago riddellii* | yes | < 0.01% | yes | 0.002% |
| *Tradescantia ohiensis* | yes | 0.01% | no | - |
| *Verbena hastata* | yes | 0.01% | no | - |
| *Vernonia fasciculata* | no | - | yes | 0.004% |
| *Zizia aurea* | yes | 0.73% | no | - |

† additional plant species included as overseed mixture in 2016

**Appendix 7. Tukey's HSD comparison of pollen diversity collected by nesting bee species**

Our best fit linear mixed model for pollen diversity within bees nests established that bee's taxonomic identity is a significant driver ($\chi^2 = 41.4$, df = 3, $p < 0.001$). Pairwise Tukey's HSD comparisons of bee species indicated that *Megachile pugnata* diet breadth was significantly greater than *Osmia caerulescens* ($p < 0.001$), *Megachile centuncularis* ($p < 0.001$), and *Osmia pumila* ($p < 0.001$). *Megachilids centuncularis* diet breadth was also greater than *O. caerulescens* ($p < 0.001$) but did not differ from *O. pumila* ($p = 0.279$). Likewise, *O. pumila* and *O. caerulescens* diet breadths were not significantly different ($p = 0.943$). Only bee species which were present in > 1 nesting straw were included in regression analyses.

| | Coef ± SE | *z* | *p* |
|---|---|---|---|
| *M. centuncularis* vs. *O. caerulescens* | 0.04 ± 0.01 | 4.62 | < 0.001* |
| *M. pugnata* vs. *O. caerulescens* | 0.13 ± 0.02 | 8.00 | < 0.001* |
| *O. pumila* vs. *O. caerulescens* | 0.02 ± 0.02 | 1.06 | 0.696 |
| *M. pugnata* vs. *M. centuncularis* | 0.09 ± 0.02 | 5.16 | < 0.001* |
| *O. pumila* vs. *M. centuncularis* | -0.03 ± 0.02 | -1.30 | 0.537 |
| *O. pumila* vs. *M. pugnata* | -0.12 ± 0.02 | -4.81 | < 0.001* |

**Appendix 8.** Proportion of identified plant genera DNA for each bee taxa collected. Status reflects whether a plant was native or exotic in Ohio, United States. Origin reflects whether a plant species corresponding to an observed pollen genus, was intentionally seeded and observed in our study habitats (*urban farm, urban prairie*), was common, urban spontaneous vegetation (Turo et al. 2021), or was not observed in our study system and grew elsewhere (residential gardens, parks, etc.). Woody notes whether a plant was a woody tree or shrub or not. Only plant genera present in ≥1.00% of total pollen collected per species are reported below.

| bee taxa | plant genus | pollen % | status | origin | woody |
|---|---|---|---|---|---|
| *Megachile campanulae* | *Lotus* | 95.57% | exotic | USV | not-woody |
| *Megachile campanulae* | *Koelreuteria* | 3.44% | exotic | Other | woody |
| *Megachile centuncularis* | *Securigera* | 18.33% | exotic | USV | not-woody |
| *Megachile centuncularis* | *Lotus* | 16.55% | exotic | USV | not-woody |
| *Megachile centuncularis* | *Cichorium* | 10.03% | exotic | USV | not-woody |
| *Megachile centuncularis* | *Taraxacum* | 7.85% | exotic | USV | not-woody |
| *Megachile centuncularis* | *Asparagus* | 6.13% | exotic | Farm | not-woody |
| *Megachile centuncularis* | *Rubus* | 4.27% | exotic | Farm | woody |
| *Megachile centuncularis* | *Sonchus* | 4.24% | exotic | USV | not-woody |
| *Megachile centuncularis* | *Erigeron* | 3.90% | native | USV | not-woody |
| *Megachile centuncularis* | *Arctium* | 3.64% | exotic | USV | not-woody |
| *Megachile centuncularis* | *Gleditsia* | 3.49% | exotic | USV | not-woody |
| *Megachile centuncularis* | *Trifolium* | 2.89% | exotic | USV | not-woody |
| *Megachile centuncularis* | *Zinnia* | 2.47% | exotic | Farm | not-woody |
| *Megachile centuncularis* | *Robinia* | 2.01% | native | Other | woody |
| *Megachile centuncularis* | *Rosa* | 1.89% | exotic | Farm | woody |
| *Megachile centuncularis* | *Verbesina* | 1.72% | exotic | Other | not-woody |
| *Megachile centuncularis* | *Tilia* | 1.53% | native | Other | woody |
| *Megachile centuncularis* | *Diplotaxis* | 1.03% | exotic | USV | not-woody |
| *Megachile pugnata* | *Cichorium* | 36.33% | exotic | USV | not-woody |
| *Megachile pugnata* | *Echinacea* | 11.95% | native | Prairie | not-woody |
| *Megachile pugnata* | *Sonchus* | 9.93% | exotic | USV | not-woody |
| *Megachile pugnata* | *Ratibida* | 8.92% | native | Prairie | not-woody |
| *Megachile pugnata* | *Centaurea* | 7.21% | exotic | Farm | not-woody |
| *Megachile pugnata* | *Rudbeckia* | 6.62% | native | Prairie | not-woody |
| *Megachile pugnata* | *Lotus* | 4.65% | exotic | USV | not-woody |
| *Megachile pugnata* | *Verbesina* | 3.61% | exotic | Other | not-woody |
| *Megachile pugnata* | *Taraxacum* | 3.56% | exotic | USV | not-woody |
| *Megachile pugnata* | *Lactuca* | 1.58% | exotic | USV | not-woody |
| *Megachile pugnata* | *Mikania* | 1.25% | exotic | USV | not-woody |
| *Osmia caerulescens* | *Trifolium* | 64.16% | exotic | USV | not-woody |
| *Osmia caerulescens* | *Securigera* | 11.08% | exotic | USV | not-woody |
| *Osmia caerulescens* | *Lotus* | 9.89% | exotic | USV | not-woody |
| *Osmia caerulescens* | *Vicia* | 3.97% | exotic | USV | not-woody |
| *Osmia caerulescens* | *Malva* | 1.53% | exotic | USV | not-woody |
| *Osmia caerulescens* | *Potentilla* | 1.34% | native | USV | not-woody |
| *Osmia caerulescens* | *Rubus* | 1.21% | exotic | Farm | woody |
| *Osmia caerulescens* | *Plantago* | 1.07% | exotic | USV | not-woody |

| | | | | | |
|---|---|---|---|---|---|
| *Osmia caerulescens* | *Cichorium* | 1.03% | exotic | USV | not-woody |
| *Osmia pumila* | *Trifolium* | 26.73% | exotic | USV | not-woody |
| *Osmia pumila* | *Taraxacum* | 17.29% | exotic | USV | not-woody |
| *Osmia pumila* | *Lonicera* | 13.79% | exotic | USV | woody |
| *Osmia pumila* | *Cercis* | 11.25% | native | Other | woody |
| *Osmia pumila* | *Malus* | 9.87% | native | Farm | woody |
| *Osmia pumila* | *Securigera* | 7.23% | exotic | USV | not-woody |
| *Osmia pumila* | *Lotus* | 4.50% | exotic | USV | not-woody |
| *Osmia pumila* | *Hypericum* | 2.83% | exotic | Other | not-woody |
| *Osmia pumila* | *Morus* | 2.73% | native | Other | woody |
| *Osmia taurus* | *Quercus* | 39.34% | native | Other | woody |
| *Osmia taurus* | *Chaenomeles* | 23.47% | exotic | Other | woody |
| *Osmia taurus* | *Taraxacum* | 18.92% | exotic | USV | not-woody |
| *Osmia taurus* | *Fraxinus* | 10.39% | native | Other | woody |
| *Osmia taurus* | *Malus* | 3.31% | native | Farm | woody |
| *Osmia taurus* | *Lotus* | 2.57% | exotic | USV | not-woody |
| *Pseudoanthidium nanum* | *Zinnia* | 53.10% | exotic | Farm | not-woody |
| *Pseudoanthidium nanum* | *Helianthus* | 30.23% | native | Farm | not-woody |
| *Pseudoanthidium nanum* | *Verbesina* | 6.36% | exotic | Other | not-woody |
| *Pseudoanthidium nanum* | *Cichorium* | 3.88% | exotic | USV | not-woody |
| *Pseudoanthidium nanum* | *Echinacea* | 3.66% | native | Prairie | not-woody |
| *Pseudoanthidium nanum* | *Mikania* | 1.89% | exotic | USV | not-woody |
